# Whole-Genome Sequencing of a *Janthinobacterium sp*. Isolated from the Patagonian Desert

**DOI:** 10.1101/2024.02.11.579814

**Authors:** Nicole T Cavanaugh, Girish Kumar, Alicyn Reverdy Pearson, Julia Colbert, Carlos Riquelme, André O Hudson, Yunrong Chai, Veronica Godoy-Carter

## Abstract

*Janthinobacterium* is a genus of Gram-negative environmental bacteria that survive extreme conditions by forming biofilms and producing pigments. *Janthinobacterium* sp. LS2A, an extremophile isolated from soil in the Chilean Patagonia, contains seven known biosynthetic gene clusters, including the purple pigment violacein, which may aid in its survival in harsh environments.

## Announcement

*Janthinobacterium* is a genus of Gram-negative bacteria that is widespread in cold climate soil and water (1, 2, 3). Cells of *Janthinobacterium* are rod-shaped and produce a purple pigment called violacein (4). Violacein is of high biotechnological significance due to its known antiviral, antibacterial, antimycotic, antitumor, algicidal, and antioxidant activities (5, 6, 7, 8). The genus also contains extremophiles that are tolerant to cold temperatures 2-28 ºC and UV exposure (9, 10, 11). *Janthinobacterium sp*. are known to form biofilms which help them survive these harsh conditions (4).

The goal of this project was to study extremophiles in Patagonia, Chile and the mechanisms that help them survive in an extreme environment, including pigment production and biofilm formation. Bacteria were isolated after plating 100 μL of a soil/water slurry from Laguna Sarmiento on Reasoner’s 2A (R2A) agar medium plates followed by incubation at 28 ºC for 48-72 hours. To test biofilm formation, LS2A was grown at 30 ºC to OD600 = 1.0 in R2A broth and 2 μL of culture was spotted on a dry R2A plate, which was incubated for 96 hours before imaging (Figure 1A). As a first approximation to identify closely related species, we performed 16S rRNA gene sequencing on the V3/4 variable regions using the following primers: 5’-CCTACGGGNGGCWGCAG-3’ and 5’-GACTACHVGGGTATCTAATCC-3’. The isolate was determined to be *Janthinobacterium sp*. using National Society for Biotechnology Information (NCBI) BLASTN version 2.15.0 (12, 13). The sequence of the V3/4 16S rRNA region was used, along with a variety of 16S sequences from *Janthinobacterium* sp. that were present in the RefSeq database, to create a multiple sequence alignment and a guide tree using ClustalOmega version 1.2.4 (Figure 1B) (14).

**Figure 1:**
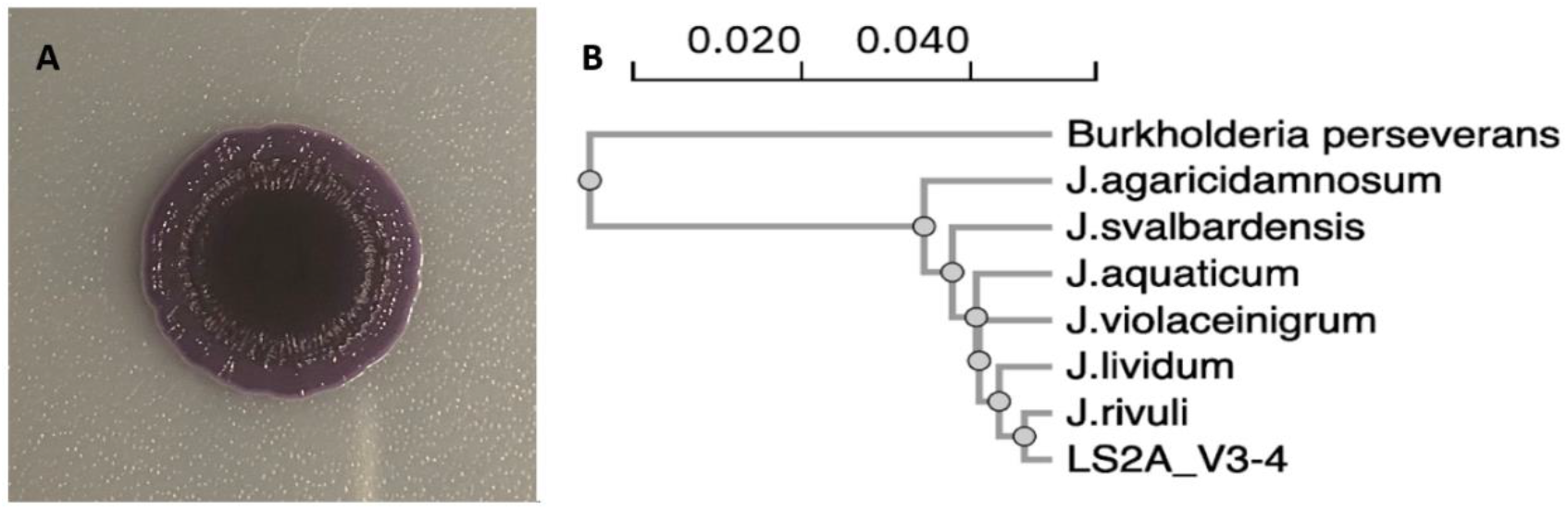
Janthinobacterium sp. LS2A forms biofilms and its sequence demonstrates it belongs to the genus. 1a. Biofilm grown from Janthinobacterium sp. LS2A. 1b. Phylogenetic Tree. A phylogenetic tree relating the LS2A V3/4 16S rRNA regions to other Janthinobacterium sp. in the RefSeq database. From top to bottom: Burkholderia preserverans NR_179094.1, J. agaricidamonosum NR_114134.1, J. svalbardensis NR_132608.1, J. aquaticum NR_170539.1, J. violaceinigrum NR_170541.1, J. lividum NR_026365.1, J.rivuli NR_170540.1, and Janthinobacterium sp. LS2A.

Genomic DNA was isolated from a 2-mL R2A liquid culture using the Wizard Genomic DNA Purification Kit (Promega, USA) following manufacturer’s instructions for Gram-negative bacteria. After quantification using a NanoDrop spectrophotometer, DNA libraries were prepared using the Nextera XT library preparation kit (Illumina) and sequenced using the Illumina MiSeq System at the Genomics Lab, Rochester Institute of Technology. Raw reads were trimmed using Trimmomatic Galaxy version 0.38.1 (15). The trimmed reads were assembled *de novo* using Unicycler Galaxy version 0.5.0+galaxy1 (16). Quality analysis was performed using Quast Galaxy version 5.2.0+galaxy1 (17) and FastQC Galaxy version 0.74+galaxy0 (18). All programs listed above were run using default parameters unless stated otherwise. The assembly consists of 4,575,745 reads assembled into 29 contigs with a read length of 6,251,303 base pairs (bp) containing 667.6 Mbp total. The N50 is 540,701 bp and the GC content is 62.69%. Analysis by the Prokaryotic Genome Analysis Pipeline (PGAP) version 6.6 revealed that the LS2A genome contained 5,664 total genes, 5,548 protein coding sequences, 3 rRNAs operons, and 77 tRNAs (19-21).

The genome assembly was analyzed using antibiotics and secondary metabolite analytics shell (antiSMASH) version 7.1.0 (22). A summary of the biosynthetic gene clusters (BGC’s) for putative antibiotics and secondary metabolites predicted by the software are outlined in Table 1. antiSMASH detected seven gene clusters that encode unique metabolites, including violacein.

**Table 1:** Predicted BGC’s in LS2A. A summary of biosynthetic gene clusters in Janthinobacterium sp. LS2A identified by antiSMASH.

| Region | Type | Start | End | Most Similar To | Similarity Score |
| --- | --- | --- | --- | --- | --- |
| 1.1 | RiPP-like | 364,750 | 372,021 | burkholderic acid | low (<10%) |
| 1.2 | Indole | 533,201 | 556,218 | violacein | 100% |
| 1.3 | RiPP-like | 830,949 | 841,878 |  |  |
| 1.4 | Acyl Amino Acids | 936,131 | 1,031,826 | O-antigen | 14% |
| 1.5 | Terpene | 1,087,968 | 1,109,729 |  |  |
| 3.1 | RiPP-like | 254,757 | 266,376 |  |  |
| 7.1 | Arylpolyene | 1 | 32,414 | APE Ec | 36% |

## Data Availability

The WGS projects for Janthinobacterium sp. LS2A have been deposited in GenBank under BioProject number PRJNA1071647, BioSample number SAMN39706722, and SRA number [SRR27839980](https://www.ncbi.nlm.nih.gov/sra/?term=PRJNA1071647). Results from PGAP analysis can be found using accession number JAZHPB000000. The following sequences were used to construct Figure 1 and were retrieved from [GenBank](https://www.ncbi.nlm.nih.gov/nucleotide/): NR_179094.1, NR_114134.1, NR_132608.1, NR_170539.1, NR_170541.1, NR_026365.1, NR_170540.1,

The [NCBI BLASTN](https://blast.ncbi.nlm.nih.gov/Blast.cgi?PROGRAM=blastn&BLAST_SPEC=GeoBlast&PAGE_TYPE=BlastSearch) webserver was used to identify the LS2A 16s rRNA gene sequence. All programs found on the Galaxy platform can be found at usegalaxy.org. [Clustal Omega](https://www.ebi.ac.uk/jdispatcher/msa/clustalo) can be found on the [EMBL European Bioinformatics Institute](https://www.ebi.ac.uk/) webpage.

## Acknowledgements

We would like to thank the National Science Foundation (MCB1651732) and the NU Global Experience Office for their support. Thank you to the RIT Genomics Lab. V.G. was supported by an HHMI Inclusive Excellence grant and the Northeastern Global Experiential Office. J.C. was supported by NU Undergraduate Research and Fellowships Office PEAK Awards. N.C. was supported by the NSF Graduate Research Fellowship Program (1938052). A.R.P. was supported by the Northeastern University Provost Dissertation Completion Fellowship.

